# PCSK9 affects expression of key surface proteins in human pancreatic beta cells through intra- and extracellular regulatory circuits

**DOI:** 10.1101/2021.09.28.461975

**Authors:** Kevin Saitoski, Maria Ryaboshapkina, Ghaith M. Hamza, Andrew F. Jarnuczak, Claire Berthault, Françoise Carlotti, Mathieu Armanet, Kaushik Sengupta, Christina Rye Underwood, Shalini Andersson, Isabelle Guillas, Wilfried Le Goff, Raphael Scharfmann

## Abstract

**Aims/hypothesis:** Proprotein convertase subtilisin/kexin 9 (PCSK9) is involved in the degradation of LDLR. However, PCSK9 can target other proteins in a cell-type specific manner. While PCSK9 has been detected in pancreatic islets, its expression in insulin-producing pancreatic beta cells is debated. Herein, we studied PCSK9 expression, regulation and function in the human pancreatic beta cell line EndoC-βH1.

**Methods:** We assessed PCSK9 expression in mouse and human pancreatic islets, and in the pancreatic beta cell line EndoC-βH1. We also studied PCSK9 regulation by cholesterol, lipoproteins, Mevastatin, and by SREBPs transcription factors. To evaluate PCSK9 function in pancreatic beta cells, we performed PCSK9 gain-and loss-of-function experiments in EndoC-βH1 using siPCSK9 or recombinant PCSK9 treatments, respectively.

**Results:** We demonstrate that PCSK9 is expressed and secreted by pancreatic beta cells. In EndoC-βH1 cells, PCSK9 expression is regulated by cholesterol and by SREBPs transcription factors. Importantly, PCSK9 knockdown results in multiple transcriptome, proteome and secretome deregulations and impaired insulin secretion. By gain- and loss-of-function experiments, we observed that PCSK9 regulates the expression levels of LDLR and VLDLR through an extracellular mechanism while CD36, PD-L1 and HLA-ABC are regulated through an intracellular mechanism.

**Conclusions/interpretation:** Collectively, these results highlight PCSK9 as an important regulator of CD36, PD-L1 and HLA-ABC cell surface expression in pancreatic beta cells.

**Data availability:** RNA-seq data have been deposited to GEO database with accession number GSE182016. Mass spectrometry proteomics data have been deposited to the ProteomeXchange Consortium via the PRIDE partner repository with the following identifiers: PXD027921, PXD027911 and PXD027913.

## Introduction

Proprotein convertase subtilisin/kexin type 9 (PCSK9) is the last discovered member of the proconvertases family [1]. It is mainly expressed in liver, cerebellum, lung and organs of the gastro-intestinal tract including pancreas (https://gtexportal.org/home/gene/PCSK9) [2]. Hepatocytes express the highest levels of PCSK9 and are the source of circulating PCSK9 [3]. PCSK9 is synthesized as an immature zymogen precursor and is autocatalytically cleaved in the endoplasmic reticulum between its N-terminal pro-domain and catalytic domain [4]. The resulting mature PCSK9 is next released into the circulation where it plays a key role in the regulation of cholesterol homeostasis [3– 5].

PCSK9 binds extracellularly to the Low Density Lipoprotein (LDL) Receptor (LDLR) and targets it for lysosomal degradation, preventing its recycling to the plasma membrane [4]. In addition, a number of studies suggest that PCSK9 can also decrease LDLR abundance at the cell surface prior to its secretion via an intracellular route. Indeed, PCSK9 and LDLR interact early in the secretory pathway [6] and PCSK9 favors LDLR degradation even when its endocytosis is blocked [7]. Moreover, inhibition of LDLR traffic between the trans Golgi network and the lysosome abolishes PCSK9 induced LDLR degradation [8].

In addition to LDLR, PCSK9 enhances the degradation of other proteins: Very Low Density Lipoprotein Receptor (VLDLR), LDL-related protein-8 (LRP8 also called ApoER2) [9], Cluster of differentiation 81 (CD81) [10], LDL-related protein-1 (LRP1) [11], Cluster of differentiation 36 (CD36) [12], β-secretase 1 (BACE1) [13] as well as the epithelial sodium channel subunits (SCNN1A, SCNN1B and SCNN1G, also called α-, β- and γ-ENaC) [14]. Very recently, it was also demonstrated that PCSK9 disrupts the recycling of major histocompatibility complex (MHC I) to the cell surface, thus positioning PCSK9 inhibitors as a new way to enhance immune checkpoint therapy for cancer [15]. Some of the above-described proteins including the LDLR have been shown to be regulated by PCSK9 both intracellularly and extracellularly [9–12, 15]. For other proteins such as BACE1, intracellular PCSK9-mediated protein degradation has been described as an important route [13]. Moreover, it is the unique route for the epithelial sodium channel ENaC degradation by PCSK9 [14]. Finally, for other proteins, their route of degradation by PCSK9 is far less studied.

Regulation of PCSK9 expression in the liver and hepatocytes has been deeply studied [4]. A Sterol Regulatory Element (SRE) has been identified in the PCSK9 promotor [16] and the Sterol Regulatory Element-Binding Protein (SREBP) transcription factors play a major role in this regulation [17, 18]. In hepatocytes, PCSK9 expression is dependent on both SREBP-1 and SREBP-2 proteins, while the regulation of PCSK9 by cholesterol is predominantly SREBP-2 dependent [16–18]. PCSK9 expression is also modulated by insulin in a SREBP-1c dependent manner [19].

PCSK9 gain-of-function mutations are associated with familial hypercholesterolemia [20] while patients with loss-of-function PCSK9 mutations have reduced circulating LDL-cholesterol levels [21]. These discoveries led to validation of PCSK9 as a therapeutic target for the treatment of hypercholesterolemia. The first anti-PCSK9 monoclonal antibodies were approved for the treatment of hypercholesterolemia in 2015 [22] and recently, small interfering RNAs targeting intracellular PCSK9 were approved by the European Medicines Agency (EMA) for the treatment of hypercholesterolemia or mixed dyslipidemia [23]. Interestingly, PCSK9 loss-of-function mutations are associated with increased prevalence of type 2 diabetes [24]. There is no evidence of an effect of anti-PCSK9 inhibitory antibodies on the transition to new onset diabetes, suggesting that reducing circulating PCSK9 does not impair pancreatic beta cell function [25–27].

The islets of Langerhans are micro-organs distributed throughout the pancreas. They contain beta, alpha, delta, PP and epsilon endocrine cells that produce and release insulin, glucagon, somatostatin, pancreatic polypeptide and ghrelin, respectively. PCSK9 is expressed in mouse and human pancreatic islets [28, 29]. However, the precise cell type expressing PCSK9 within islets remains unclear. Some studies demonstrate that PCSK9 is only expressed in somatostatin-producing delta cells [28, 30], while other studies indicate that beta cells express PCSK9 [29, 31, 32]. The function of PCSK9 within islets is also debated. Global Pcsk9-deficient mice were glucose intolerant and insulinopenic [29, 30], though, different studies showed conflicting results [28, 31]. These discrepancies might be due to differences in the genetic background and age of the mice. Finally, PCSK9 function in islets has only been studied through the regulation of LDLR [28–30] without a search for additional targets.

The aim of this study was to clarify the role of PCSK9 in human pancreatic beta cells. By using the human pancreatic beta cell line EndoC-βH1 [33], we investigated PCSK9 expression, function and its regulation. We also searched for additional cell surface proteins regulated by PCSK9 in human pancreatic cells by performing gain- and loss-of-function experiments.

## Experimental procedure

### Ethical statement

This study was performed according to the Declaration of Helsinki and the Declaration of Istanbul. No tissues were procured from prisoners. Human islets were provided by the Leiden University Medical Center (Leiden, The Netherlands) and the human islet core facility of St-Louis Hospital (Paris, France). They were prepared from pancreata of brain-dead donors after informed consent was signed from next-of-kin, and processed for islet isolation according to the procedures approved by the ethics committee of the French Agency of Biomedicine, and the Dutch national law. All of the animal studies complied with the ARRIVE guidelines and were conducted in strict accordance with the EU Directive 2010/63/EU for animal experiments and with regard to specific national laws and INSERM guidelines.

### Culture of EndoC-βH1 cells and treatment

The human pancreatic beta cell line EndoC-βH1 cells (Univercell Biosolutions, Toulouse, France [mycoplasma negative]) was cultured as previously described [33]. Experiments were carried out 24 hours after seeding the cells at a density of approximately 10^5^ cells /cm^2^. The following compounds were used for treatment: Cholesterol-Methyl-β-Cyclodextrin (Sigma-Aldrich, France), Mevastatin (Sigma-Aldrich, Germany), human native LDL and copper oxidized LDL or VLDL (see below for lipoproteins extraction protocol), Human Recombinant Wild Type PCSK9 (Circulex).

### Human hepatocytes, HepG2 and HEK293T culture

Human primary hepatocytes were isolated from liver fragments obtained from adult patients undergoing partial hepatectomy as previously described [34] and cultured in William’s Medium E (Life Technologies) supplemented with 10% FCS (Perbio), 50 µM hydrocortisone hemisuccinate (SERB laboratory), 5 μg/ml insulin (Sigma-Aldrich), 2 mM L-glutamine and 200 U/ml penicillin, 200 μg/ml streptomycin. The human hepatocellular carcinoma cell line HepG2 and the human embryonic kidney (HEK293T) (mycoplasma free) were cultured in Dulbecco’s Modified Eagle’s Medium (Gibco) and DMEM/F12 (Gibco), respectively, supplemented with 10 % FCS, 100 units/mL penicillin, and 100 μg/mL streptomycin.

### Human islets

Islets were cultured in 12-well plates in CMRL medium supplemented with 10% FCS, Hepes and penicillin/streptomycin (all from Thermo Fisher Scientific). Donor’s characteristic and islets details are presented in **Supplementary Table 1**.

### siRNA transfection

EndoC-βH1 cells were transfected as described [35] in OptiMEM using Lipofectamine RNAiMAX (Life Technologies) with siRNA SMARTpools (Horizon Discovery LTD, UK). Medium was replaced 2.5 hours later with fresh culture medium and analyses were carried out 3 to 6 days following transfection. siRNA reagents used: Control non-targeted siRNA (siCTRL, D-001810-01-20); siRNA targeting: PCSK9 (siPCSK9) (M-005989-01-0005), SREBF-1 (siSREBF-1) (M-006891-00-0005) and SREBF-2 (siSREBF-2) (M-009549-00-0005) at the final concentration of 80 nM.

### Cell sorting of mouse alpha, beta and delta cell populations

Alpha, beta and delta cell population were prepared from mouse pancreatic islets as previously described [36].

### Lipoproteins preparation and LDL oxidation

LDL and VLDL were extracted from the plasma of healthy volunteers, by sequential ultracentrifugations, as previously described [37]. Protein concentration of the lipoproteins was determined by using BCA assay (Pierce). Copper-oxidation of LDL was done by incubating the native LDL with 5 µmol/l CuSO_4_ at 37°C for 16 hours followed by dialysis (3 times) against PBS containing 0.1 mM ethylenediamine tetraacetic acid and storage at 4°C.

### RNA isolation, reverse transcription, and qPCR

A RNeasy Micro Kit (Qiagen) was used to extract total RNA from EndoC-βH1 cells [38]. Genomic DNA was removed by DNAse treatment following the RNeasy Micro Kit protocol. RNAs were reverse transcribed by using the Maxima First Strand cDNA kit (Thermo Fisher Scientific). RT-qPCR was performed using Power SYBR Green mix (Applied Biosystems) with a QuantStudio 3 analyzer (Thermo Fisher Scientific). Custom primers were designed with Primer-Blast online, and their efficiency and specificity were determined for each pair by RT-qPCR on a serial dilution of cDNA samples. The list of primers is presented in the **Supplementary Table 2**. Relative quantification (2^^-dCt^) was used to calculate the expression levels of each target gene, normalized to *CYCLOPHILIN-A* transcripts.

### Western blot analysis

Western blot experiments were performed as described [38]. For VLDLR immunoblot, cell extracts were loaded in 8 % Bis-Tris polyacrylamide gels in absence of heating and 2-beta-mercaptoethanol. The following antibodies were used: PCSK9 (1/200; AF3888; Bio-Techne), diluted in TBS 0.1 % Tween with 0,5% BSA; LDLR (1/1,000; ab52818; Abcam), VLDLR (1/1,000; ab75591; Abcam), alpha-Tubulin (1/2,000, T9026; Sigma-Aldrich), beta-actin (1/2,000; A5441; Sigma-Aldrich). Species-specific HRP-linked secondary antibodies (1/1,000; 7074 and 7076; Cell Signaling Technology) were used. Antibodies were validated by knockdown experiments (PCSK9) or or have passed application specific testing standards.

### PCSK9 secretion

EndoC-βH1, HepG2, HEK293T and human islets were starved for one hour in their respective culture media (free from BSA or FCS) and then cultivated for 24 hours in fresh BSA/FCS-free culture medium. Conditioned media were collected, centrifugated to eliminate cell debris, and used for Western blot analyses. In the case of human islets, secreted proteins were enriched following precipitation with acetone before Western blot analysis.

### Flow cytometry

EndoC-βH1 cells were trypsinized, washed three times in HBSS/1% BSA and incubated at 4 °C for 15 minutes in the dark with the following antibodies: LDLR (1/100; MAB CL 472413; Fisher scientific), CD36 (1/10, 555455, BD Pharmingen), PD-L1 (1/100, 329714, Biolegend), HLA-ABC (1/100, 3114410, Biolegend) diluted in HBSS medium/1% BSA (Gibco/Roche Diagnostics). Fluorochrome-conjugated primary antibodies were used with the exception of LDLR detection, for which a step of incubation with Alexa-fluor 488 (1/400, Invitrogen) secondary antibody was performed. Following rinsing in HBSS/1% BSA and incubation with FACS medium containing propidium iodide (1/4,000, Sigma-Aldrich), analysis was carried out using a FACS Aria III (BD Biosciences, San Jose, CA, USA). Dead cells were excluded from analyses by using propidium iodide. Data were analyzed using FlowJo 10.7 software (RRID: RRID:SCR_008520). Results are expressed in mean fluorescence intensities fold changes relative to the control condition.

### Insulin secretion

EndoC-βH1 cells were transfected with siControl (siCTRL) or siPCSK9 (siPCSK9). Two days later, cells were starved in DMEM containing 0.5 mM glucose for 24 hours. Then, cells were washed twice and then preincubated in Krebs-Ringer bicarbonate HEPES buffer (KRBH) containing 0.2% fatty acid–free BSA in absence of glucose for 1 hour. Insulin secretion was measured following a 1-hour incubation with KRBH containing 0.2% fatty acid–free BSA and 0 mM or 20 mM glucose, or 50 mM KCl. Insulin secretion and intracellular insulin were measured by ELISA as previously described [33].

### Fatty acid uptake

Fatty acid uptake in EndoC-βH1 was studied using the Fatty Acid Uptake Kit (Sigma-Aldrich). Cells were transfected with siControl (siCTRL) or siPCSK9 (siPCSK9). Six days later, cells were starved for 1 hour in cell culture medium free from BSA. TF2-C12 fluorescent fatty acid was added to the culture medium for 1 hour. Fluorescence signal was measured at λexit=485nm/λem=520nm and normalized to cell count. Fatty acid uptake was calculated by subtracting the fluorescence intensity measured at 1 hour incubation by that measured at time point zero, following the manufacturer’s instructions. Results were expressed in fatty acid uptake fold changes relative to control.

### RNA-seq and proteomics analyses

Acquisition of proteomics and RNA-seq data and sample size calculation for omics experiments have been previously described [39]. Benjamini-Yekutielli method [40] was used to adjust for multiple testing for IPA Pathway terms due to non-ignorable overlap between genes underlying top pathways. Gene set enrichment analysis was conducted with GSEA v4.0.3 software (RRID: SCR_003199).

### Statistical analysis of RT-qPCR, Western blot and FACS experiments

Each *n* represents an independent experiment. All of the graphs show means +/- Standard Error of the Mean (SEM) and the statistical analysis were conducted using Prism (GraphPad, San Diego, CA, USA) either with t-test if the normality condition was respected or with the non-parametric equivalent Mann-Whitney test. A *p-value* less than 0.05 was considered significant and Symbols for indicating *p-values* are *p < 0.05, **p < 0.01, and ***p < 0.005.

## Results

### PCSK9 is expressed and secreted by pancreatic beta cells

The cell types expressing PCSK9 in mouse and human pancreatic islets remain debated [28–31]. Here, we first analyzed *PCSK9* expression in mouse pancreatic islet cell types by using a FACS-based strategy to purify beta, alpha and delta cell populations (**Fig. S1a**) [36]. RT-qPCR analyses indicated that alpha (CD71^-^), beta (CD71^+^) and delta (CD24^+^) populations were highly enriched in *glucagon, insulin-1* and *somatostatin* mRNA, respectively, validating the efficiency of the sorting (**Fig. S1b**). *PCSK9* mRNA analyses indicated that its expression was 3-fold higher in beta than in the delta and alpha cell populations (**Fig. 1a**). PCSK9 mRNA level was also twofold lower in beta cells than in mouse liver (**Fig. 1a**). We then compared *PCSK9* mRNA expression by RT-qPCR in human islet preparations versus human primary hepatocytes and HEK293T cells (used as positive and negative controls, respectively). *PCSK9* mRNA were detected in human islets, although at a level fivefold lower than in human primary hepatocytes. As expected, *PCSK9* was almost undetectable in HEK293T cells (**Fig. 1b**). We next investigated PCSK9 expression in human islet cell types. Since PCSK9 expression in bulk islets was low, it may be difficult to capture with single-cell sequencing due to limitations of the technology. Therefore, we used a combination of scRNA-seq (E-MTAB-5061 [41], GSE81608 [42], GSE86469 [43]) and RNA-seq on highly purified FACS sorted human beta and alpha cells (GSE67543 [44]) to analyze PCSK9 expression. PCSK9 transcripts were enriched in human adult beta cells (**Fig. S1c**). We next analyzed *PCSK9* expression in the human pancreatic beta cell line EndoC-βH1 [33]. There, *PCSK9* mRNA level was higher than in the human hepatoma cell line HepG2 (**Fig. 1c**). Western blot analysis revealed the presence of two major PCSK9 bands at ∼75 kDa and ∼60 kDa. These bands corresponded to the expected sizes for pro-PCSK9 and mature PCSK9. Quantification of mature PCSK9 level showed a 3-fold higher expression in EndoC-βH1 compared to HepG2 cells, and no expression in HEK293T cells (**Fig. 1d and e**). Noteworthy, PCSK9 maturation rate (as measured by the mature PCSK9/total PCSK9 ratio) was higher in HepG2 cells than in the EndoC-βH1 (**Fig. S1d**). Given the fact that PCSK9 is a secreted protein, we also compared by Western blot PCSK9 levels in conditioned media from EndoC-βH1, HepG2 and HEK293T cells. In EndoC-βH1 and HepG2 cells, we detected a single band at approximately 60 kDa (as expected for the secreted mature form of PCSK9) and no signal in conditioned media from the HEK293T cells (**Fig. 1f**). PCSK9 secretion was twice lower in EndoC-βH1 when compared to HepG2 cells (**Fig. 1f and g**). It was also the case when PCSK9 secretion was normalized to PCSK9 cell content (**Fig. S1e**). Finally, as *PCSK9* transcripts were present in human islet preparations (**Fig. 1b**), we evaluated whether PCSK9 was secreted by human islets. By Western blot, we detected in conditioned media the presence of a band at ∼60 kDa, the expected size for the secreted form of PCSK9 (**Fig. 1h**). Omics data analysis confirmed that EndoC-βH1 cells are a representative *in vitro* model for human beta cells [39]. Thus, mouse and human beta cells express PCSK9, and EndoC-βH1 cells can be used as a model to study PCSK9 regulation and function in human beta cells.

**Figure 1:**
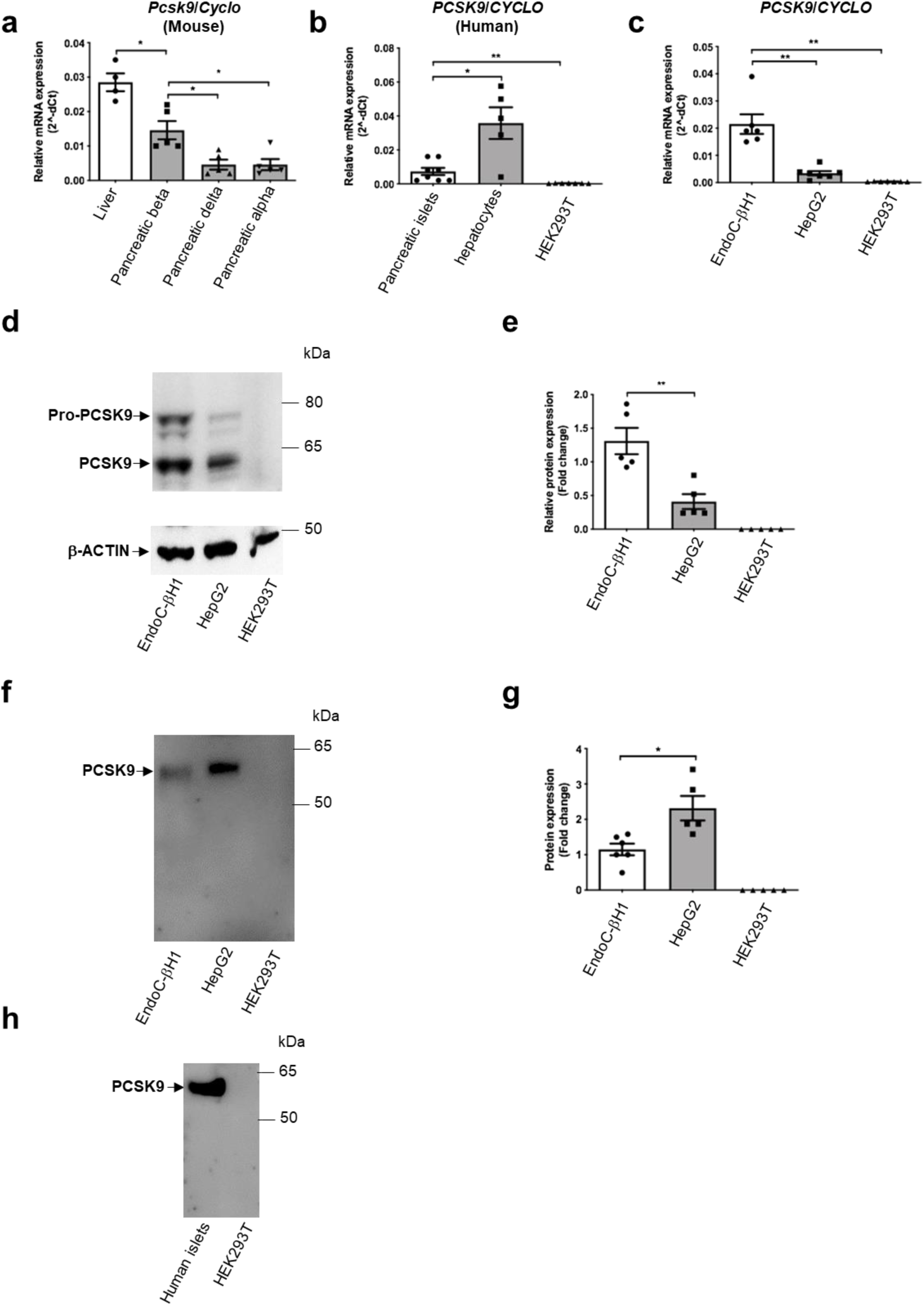
PCSK9 is expressed and secreted by pancreatic beta cells. (**a-c**) *PCSK9* mRNA expression in: (a) mouse liver compared to FACS-enriched pancreatic beta, delta and alpha endocrine fractions (*n=4-5*); (b) human islets, hepatocytes and HEK293T cells (*n=5-8*); (c) EndoC-βH1, HepG2 and HEK293T cells (*n=6-7*). **(d, e)** Detection by Western blot and quantification of intracellular PCSK9 in EndoC-βH1, HepG2 and HEK293T cells (*n=5*). (**f, g**) Detection by Western blot and quantification of secreted PCSK9 in EndoC-βH1, HepG2 and HEK293T cells (*n=5-6*). (**h**) Western blot analysis of PCSK9 secretion by human islets, compared to HEK293T cells (*n=7*). Data represent the means ± SEM. *p < 0.05, **p < 0.01

### Cholesterol, LDL and mevastatin modulate PCSK9 expression and secretion

In mouse and human hepatocytes, PCSK9 expression is regulated by cholesterol and by lipid-lowering drugs of the statin class [16, 45]. Here, we evaluated the effects on PCSK9 expression of cholesterol, LDL and mevastatin in EndoC-βH1 cells. Cholesterol and LDL treatments reduced the mRNA expression of both SREBP2 (*HMGCR* and *LDLR)* and SREBP1 (*FASN* and *SCD*) transcriptional targets by ∼50%. Mevastatin treatment increased the expression of these genes by at least two-fold, demonstrating the functionality of the LDL-cholesterol pathway in EndoC-βH1 cells (**Fig. 2a**). *PCSK9* mRNA followed a similar pattern as expression was reduced by cholesterol and LDL treatment and increased following incubation with Mevastatin (**Fig. 2b)**. The effects of Mevastatin were reverted by cholesterol or LDL treatments and were thus “cholesterol specific “(**Fig. 2a, b**). Similar effects of cholesterol, LDL and Mevastatin were also observed at protein level. Indeed, Western blot analyses indicated that cholesterol and LDL treatments decreased, while Mevastatin increased intracellular PCSK9 levels (**Fig. 2c and d**). Similar results were also obtained when secreted PCSK9 was analyzed (**Fig. 2e and f**). Finally, we tested whether other classes of human lipoproteins modulate *PCSK9* expression in EndoC-βH1 cells. Treatment with Triglyceride-Rich Lipoproteins (TRL) mimicked the effects of *LDL* on *PCSK9* mRNA expression and on *SREBP-1* and *SREBP-2* transcriptional targets while oxidized LDL had no effect (**Fig. 2g, h**).

**Figure 2:**
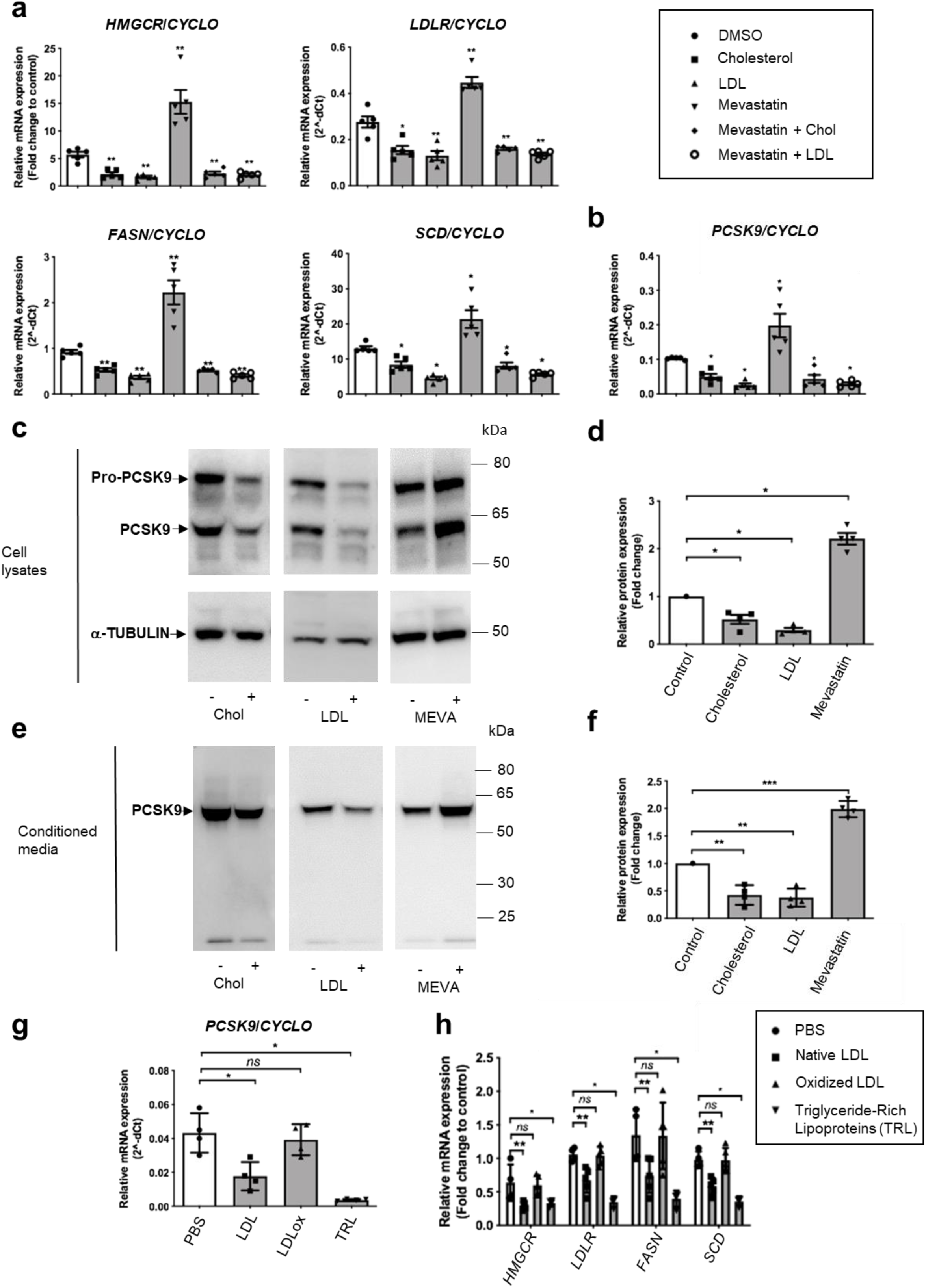
Cholesterol, LDL and mevastatin modulate PCSK9 expression and secretion. (**a-f**) EndoC-βH1 cells were exposed to the indicated treatments for 24 hours. (a, b) SREBP-2 transcriptional targets (*HMGCR* and *LDLR*), SREBP-1 transcriptional targets (*FASN*, and *SCD*) and PCSK9 mRNA were measured by RT-qPCR (*n=5*). Intracellular (c, d) and secreted (e, f) PCSK9 were analyzed and quantified by Western blot (*n=4*). (**g, h**) EndoC-βH1 cells were exposed to native LDL, oxidized LDL or Triglyceride-Rich Lipoproteins (TRL) for 24 hours. *PCSK9*, SREBP1 and SREBP2 targets were measured by RT-qPCR. (*n=4*) Data represent the means ± SEM. *p < 0.05, **p < 0.01 and ***p < 0.001

### PCSK9 is regulated by the SREBPs-1 and 2 transcription factors

We first tested whether *PCSK9* mRNA levels were dependent on SREBPs transcription factors. Co-transfection of the EndoC-βH1 cells with siSREBF-1+siSREBF-2 efficiently reduced mRNA expression of *SREBF-1, SREBF-2* and their respective target genes: *FASN, SCD* (SREBP-1 targets) and *HMGCR, LDLR* (SREBP-2 targets). *PCSK9* mRNA levels were downregulated indicating dependence of SREBPs activity (**Fig. 3a**). Single siSREBF-2 downregulated *SREBF-2* and its target *HMGCR*, but not *SREBF-1*. It increased PCSK9 mRNA levels (**Fig. 3b**). Single siSREBF-1 transfection specifically reduced *SREBF-1* expression. The expression of canonic SREBP-1 liver targets *FASN* and *SCD* was not modulated, indicating the existence of compensatory mechanism that maintain *FASN, SCD* and *LDLR* expression when SREBF-1 or SREBF-2 are reduced following knockdown (**compare Fig. 3a to 3b and 3c**). On the other hand, *SREBF-1* knockdown was sufficient to decrease *PCSK9* expression, indicating that SREBF-1 is essential to maintain PCSK9 expression (**Fig. 3c**).

**Figure 3:**
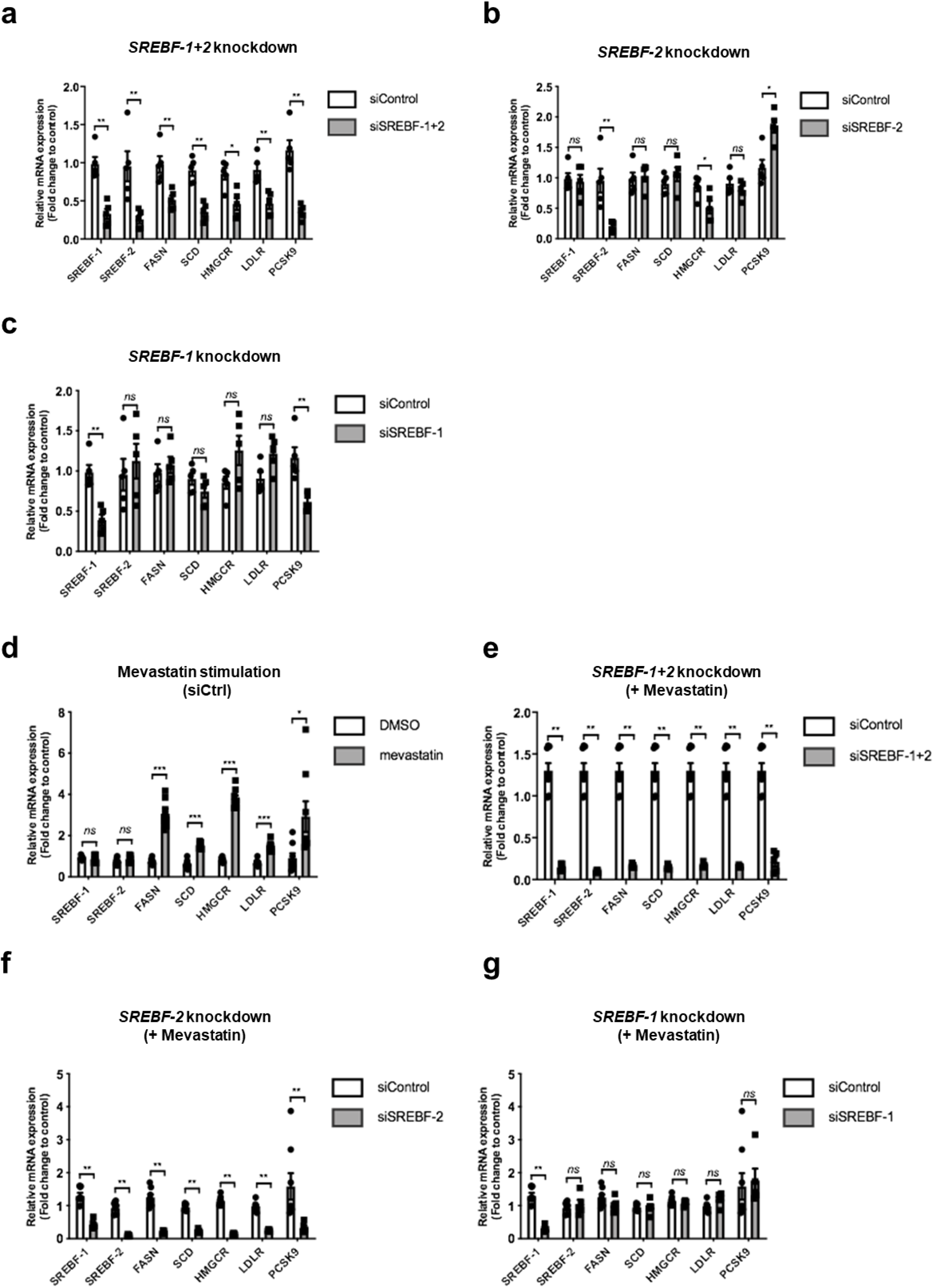
PCSK9 is regulated by the SREBPs-1 and 2 transcription factors. (**a-c**) EndoC-βH1 cells were transfected with control nontarget siRNA (siCtrl), siRNA targeting SREBP-1+SREBP-2 (siSREBP-1+2), SREBP-2 (siSREBP-2), or SREBP-1 (siSREBP-1). RT-qPCR for SREBP’s and their targets were performed 3 days later (*n=5*). (**d-g**) EndoC-βH1 cells were transfected with siRNA, cultured for 72h and next treated for 24 hours with DMSO (control) or mevastatin before analyses by RT-qPCR (*n=5*). Data represent the means ± SEM. *p < 0.05, **p < 0.01 and ***p < 0.001

We next assessed the SREBPs dependence of PCSK9 upon Mevastatin treatment. As expected, Mevastatin treatment increased the mRNA expression of SREBP-1 targets, SREBP-2 targets and also *PCSK9* (**Fig. 3d**). This induction was reverted upon SREBF-1+2 double knockdown (**Fig. 3e**). Single SREBF-2 knockdown reduced the expression of *SREBF-2* and its target genes *HMGCR* and *LDLR*. PCSK9 expression was also down-regulated suggesting a SREBP-2 dependence (**Fig. 3f**). However, we noted that upon Mevastatin treatment, siSREBF-2 decreased *SREBF-1* levels (**Fig. 3f**). We thus performed single siSREBF1 knockdown that did not modulate *PCSK9* expression (**Fig. 3g**). Taken together, our data indicate that SREBP-2, but not SREBP-1 is involved in the induction of PCSK9 by Mevastatin in human beta cells.

### Insights from hypothesis-generating omics analysis

We investigated the effect of siRNA-mediated PCSK9 knockdown in EndoC-βH1 cells in transcriptomics and proteomics experiments. PCSK9 knockdown was confirmed in all omics experiments (**Fig. S2**). Changes in gene and protein expression levels were generally consistent and had the same directionality (60.3% for RNA-seq vs DIA cell lysate proteomics and 59.7% for RNA-seq vs TMT cell lysate proteomics) (**Fig. S3**). IPA Pathway enrichment analysis indicated impaired mitochondrial function (**Fig. 4a**) which was confirmed by GSEA analysis (**Fig 4b**). Normal mitochondrial function is crucial for insulin secretion by beta cells [46]. We investigated the expression of markers associated with glucose-stimulated insulin secretion [47]. Out of 147 markers, 45 (31%) were differentially expressed **(Fig. 4c**). We also observed consistent up-regulation of PCSK1 levels, an enzyme involved in pro-insulin to insulin processing, in the transcriptome and proteome experiments (**Fig. S4**). Interestingly, we observed an up-regulation of insulin in DIA secretome proteomic at 72 hours after PCSK9 knockdown (**Fig. S4a**). Following PCSK9 knockdown, insulin secretion was impaired: basal and glucose-stimulated insulin secretion increased, while KCl-induced insulin secretion was reduced **(Fig.4d)**. Gene Ontology Cellular Component enrichment analysis indicated overrepresentation of MHC-I complex and plasma membrane proteins among genes and proteins up-regulated after PCSK9 knockdown (**Fig. 4e**). MHC I complex enrichment was stronger on protein level than on mRNA level. Golgi and endoplasmic reticulum proteins were also up-regulated, which is consistent with the reported role of PCSK9 in intracellular degradation of proteins in the ER-to-Golgi intermediate compartment [13]. CD36, CD274 (PD-L1) and MHC I complex proteins (particularly HLA-A, B, C and E) were top findings in proteomics experiments **(Fig. 4f, g, h)**. ENaC channel subunits were not expressed in Endoc-βH1 cells. LDLR, VLDL, LRP1 and LRP8 levels were not altered (**Fig. S5**). CD81 was up-regulated on mRNA but not protein level (**Fig. S5**). mRNA and whole-cell protein expression of APP and APLP2 was increased after PCSK9 knockdown (**Fig. S5**). Up-regulation of APP and APLP2 mRNA was confirmed by RT-qPCR together with the other top differentially expressed genes (**Fig. 4i**)

**Figure 4:**
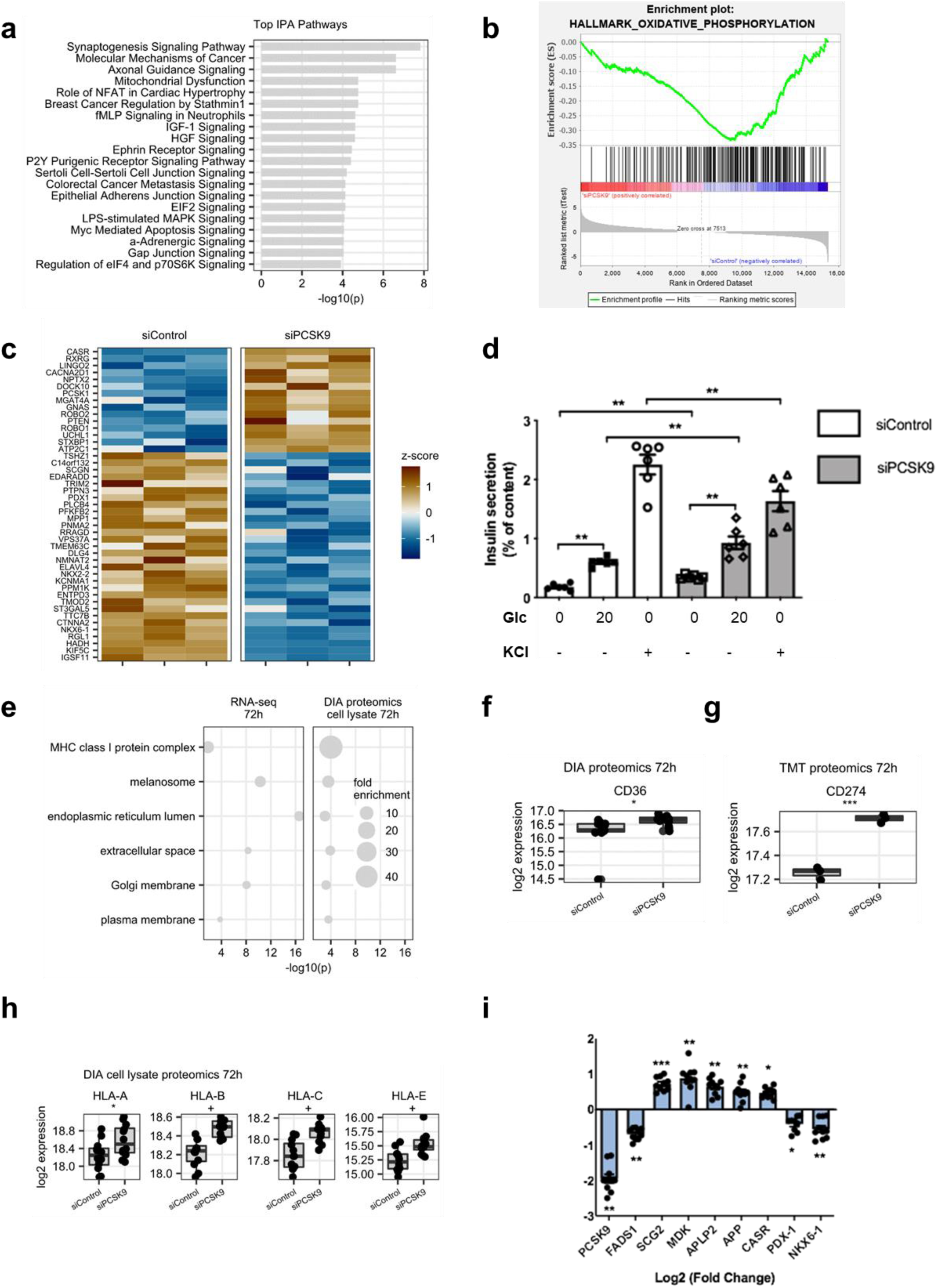
Insights from omics experiments. EndoC-βH1 cells were transfected with control nontarget siRNA (siControl) or siRNA targeting PCSK9 (siPCSK9). Three days later, cells were profiled by RNA-seq and proteomics. RNA-seq data were analyzed with: (**a**) Ingenuity Pathway Analysis (IPA) software, for the top enriched canonical pathways, and (**b**) Gene Set Enrichment Analysis (GSEA) software with the Hallmark_Oxidative_Phosphorylation gene set. (**C**) GSIS markers deregulated upon PCSK9 knockdown with FDR <0.05 in RNA-seq experiment. (**d**) Basal (0.5 mM), glucose-stimulated (20 mM) and KCl-stimulated (50 mM) insulin secretion was measured in EndoC-βH1 cells, three days following PCSK9 knockdown (*n=6*). (**e**) Gene Ontology analysis of cellular components terms enriched among genes and proteins up-regulated following PCSK9 knockdown. (**f-h**) Boxplot showing protein expression of top candidate PCSK9 degradation targets, measured on cell protein extracts by DIA and TMT proteomic. (**i**) RT-qPCR validation of selected top differentially expressed genes (*n=10*). For GSIS and RT-qPCR analyses, data are presented as means ± SEM. For proteomics experiments, median expression and interquartile ranges are shown. + FDR < 0.05, *p < 0.05, **p < 0.01 and ***p < 0.001

### PCSK9 knockdown does not affect LDLR levels

In hepatocytes, PCSK9 interacts with LDLR and favors its lysosomal degradation [4, 5, 48]. Lack of LDLR dysregulation after PCSK9 knockdown in the omics experiments were unexpected, so we evaluated the effects of exogenously added human recombinant PCSK9 on LDLR expression. Incubating EndoC-βH1 cells with recombinant PCSK9 reduced LDLR protein levels (**Fig. 5a and b**). We also performed a confirmatory siRNA experiment to test whether endogenously produced PCSK9 regulated LDLR expression. PCSK9 knockdown was efficient with a ∼80% decrease at the mRNA level (**Fig. 5c**), at the intracellular protein level (for both Pro-PCSK9 and mature PCSK9) (**Fig. 5d, f**) and at the secreted protein level (**Fig. 5e, f**). We next looked at LDLR levels. As expected, PCSK9 knockdown did not modulate LDLR mRNA levels (**Fig. 5g)**. Moreover, despite the sharp decrease in PCSK9 protein expression and secretion, PCSK9 knockdown did not impair LDLR protein levels as measured by Western blot (**Fig. 5h and i**). To determine whether PCSK9 specifically modulates LDLR at the cell surface, we performed FACS analyses. Again, PCSK9 knockdown did not modulate cell surface LDLR levels (**Fig. 5j, k**). Taken together, these results indicate that LDLR can be targeted by extracellular PCSK9 but is not regulated by intracellular PCSK9 in EndoC-βH1 cells.

**Figure 5:**
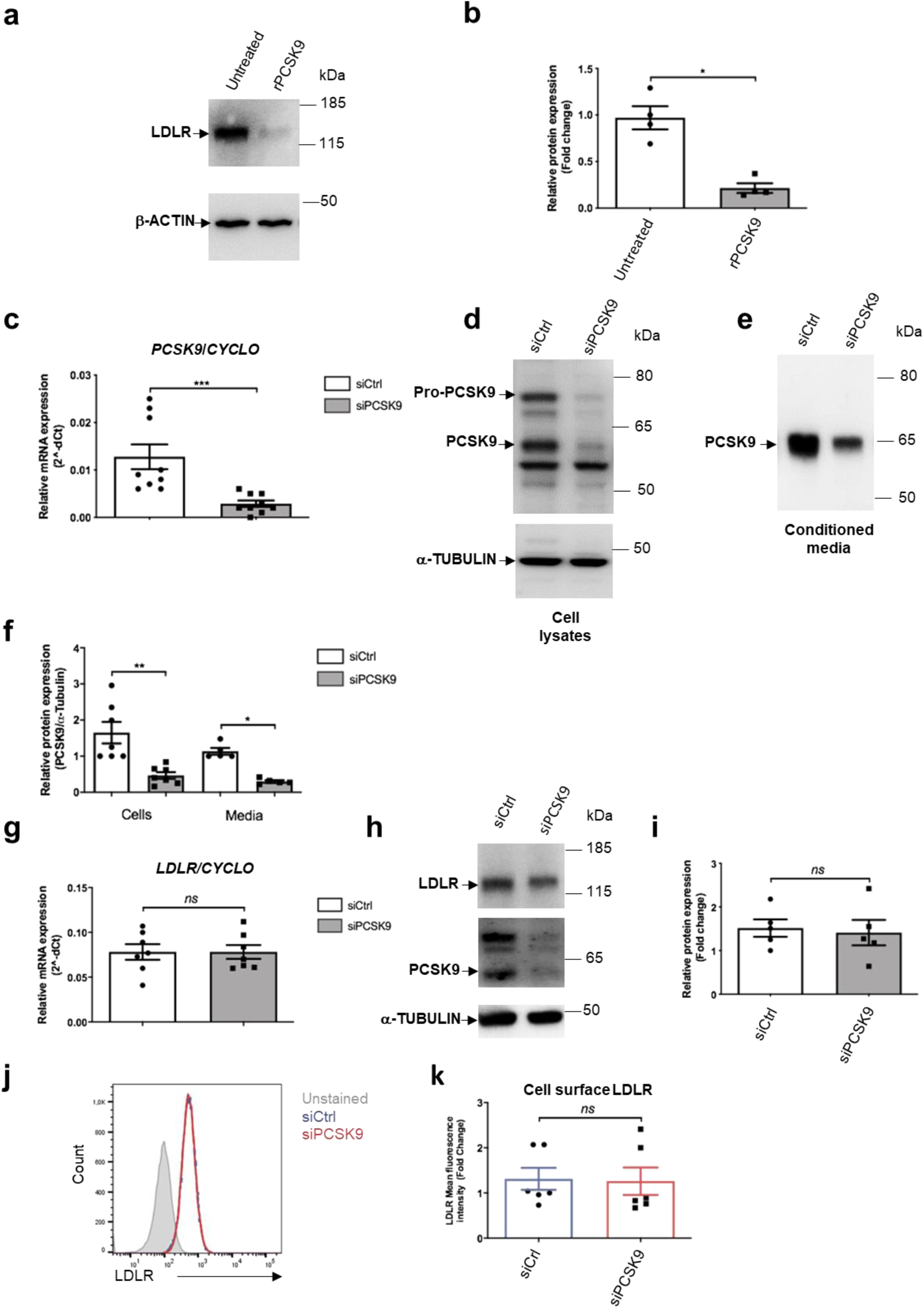
PCSK9 knockdown does not affect LDLR levels. (**a, b**): EndoC-βH1 cells were exposed for 16 hours to human recombinant PCSK9 (2.5 µg/ml). LDLR protein expression was studied by Western blot and quantified (*n=4*). (**c-k**) EndoC-βH1 cells were transfected with control nontarget siRNA (siCtrl) or siRNA targeting PCSK9 (siPCSK9). Analyses were performed 3 days later. (c-f) PCSK9 knockdown efficiency was analyzed by RT-qPCR (*n=9*) (c) and Western blot for intracellular and secreted PCSK9 (*n=5-7*) (d-f). (g-k) The effect of PCSK9 knockdown on LDLR was measured by RT-qPCR, (*n=7*), Western blot (n=5) and by FACS for cell surface expression (*n=6*). Data represent the means ± SEM. *p < 0.05, **p < 0.01 and ***p < 0.001

### PCSK9 knockdown potentiates the induction of PD-L1 and HLA-ABC by interferon-*γ*

MHC-I and PD-L1 are increased in pancreatic islets from type I diabetics and in response to Interferon class I stimulation [49–51]. While characterizing the effect of PCSK9 knockdown on the proteome of EndoC-βH1, we observed up-regulation of MHC-I (particularly HLA-A, B, C and E) and PD-L1 (**Fig. 4f, g**). We thus asked whether PCSK9 regulates PD-L1 and/or HLA-ABC cell surface expression in basal conditions and following interferon-*γ* treatment. PCSK9 knockdown consistently reduced *PCSK9* mRNA levels in basal or Interferon-*γ* stimulated conditions (**Fig. 6a**) with a slight increase of *MHC-I Mrna* under basal conditions (**Fig. 6b**). On the other hand, upon Interferon-*γ* treatment, *MHC-I* and *PD-L1* mRNA levels were unchanged following PCSK9 knockdown (**Fig. 6b**). We next measured by FACS cell surface expression of PD-L1 following siRNA-mediated PCSK9 knockdown or following treatment with recombinant PCSK9. It revealed that while siRNA-mediated PCSK9 knockdown increased cell surface expression of PD-L1 (**Fig. 6c, d**), recombinant PCSK9 did not have any effects (**Fig. 6e, f**). We performed a similar experiment, measuring cell surface expression of HLA-ABC. Again, while siRNA-mediated PCSK9 knockdown increased cell surface expression of HLA-ABC (**Fig. 6g, h**), recombinant PCSK9 did not have any effects (**Fig. 6i, j**). This data indicates that PD-L1 and HLA-ABC cell surface expression is regulated by intracellular PCSK9.

**Figure 6:**
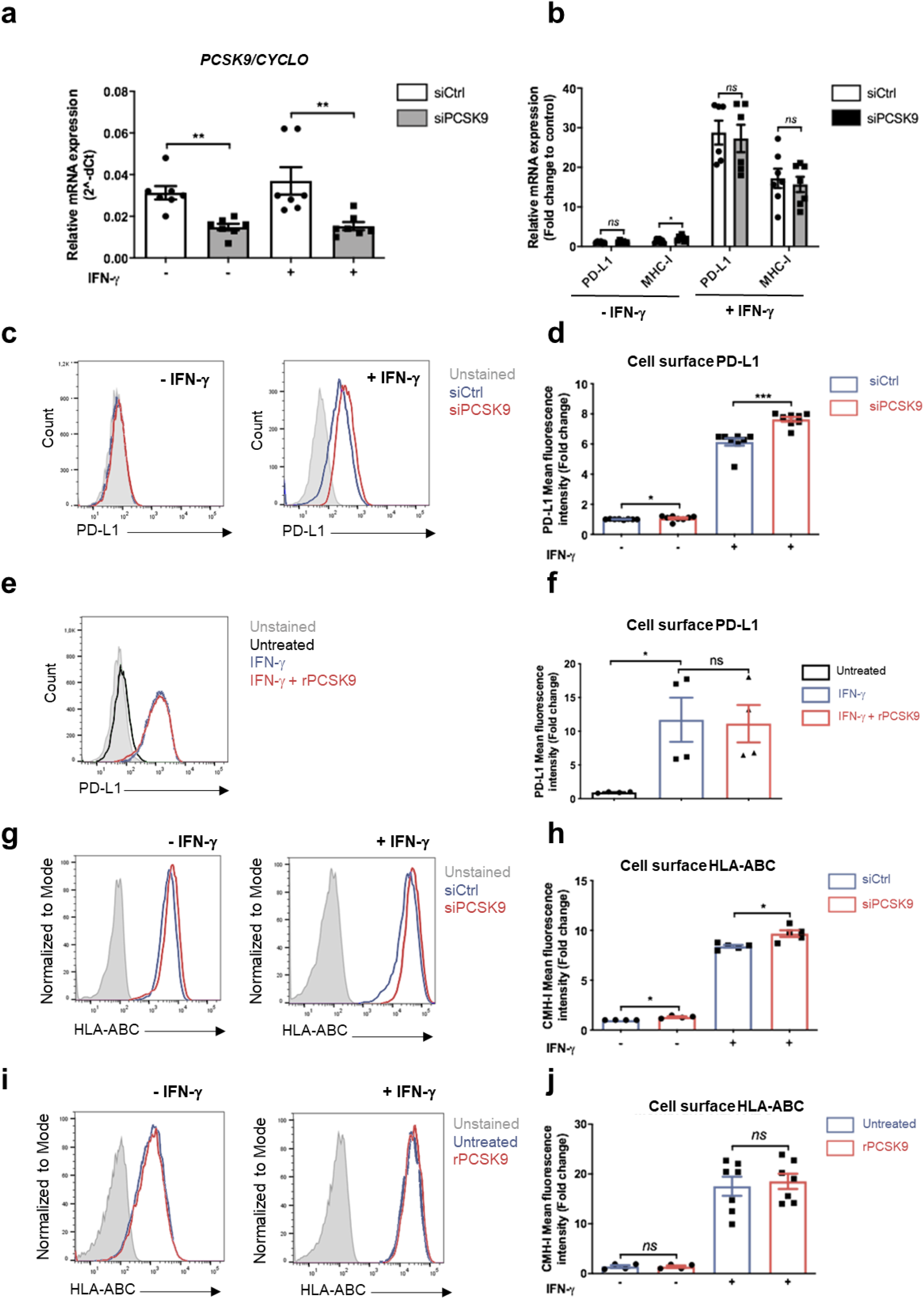
PCSK9 knockdown potentiates the induction of PD-L1 and HLA-ABC by interferon-*γ*. (**a-d**) EndoC-βH1 cells were transfected with control nontarget siRNA (siCtrl) or siRNA targeting PCSK9 (siPCSK9). Five days later, the cells were incubated for 24 hours with interferon-*γ* (IFN-*γ*). (a, b), RT-qPCR analysis of PCSK9, PD-L1 and MHC-I (*n=6-7*) (c, d) Cell surface PD-L1 expression analyzed by FACS (*n=8-9*). (**e, f**) EndoC-βH1 cells were treated with Interferon-*γ* (IFN-*γ*). Twenty-four hours later, human recombinant PCSK9 (rPCSK9) (2.5 µg/ml) was added to the culture medium for 16 hours. Cell surface PD-L1 was studied by FACS (*n=4*). (**g-j**) Cell surface HLA-ABC expression analyzed by FACS under the same experimental conditions as in (a-f) (*n=4-7*). Data represent the means ± SEM. *p < 0.05, **p < 0.01

### PCSK9 regulates the expression of VLDLR and CD36, 2 actors involved in fatty acid homeostasis

Fatty acid homeostasis is important for proper beta cell function (Lytrivi et al., 2020). Interestingly, VLDLR and CD36 are involved in the uptake of triglyceride-rich lipoproteins and fatty acids, respectively, and are PCSK9 degradation targets in liver cells [9, 12]. We further investigated their expression by RT-qPCR and Western blot. PCSK9 knockdown slightly decreased VLDLR mRNA (**Fig. S6a**) but did not impact VLDLR protein expression (**Fig. 7a, b**). Incubating EndoC-βH1 cells with recombinant PCSK9 reduced VLDLR protein expression (**Fig. 7c, d)**. PCSK9 knockdown cells exhibited a clear reduction in PCSK9 mRNA level (**Fig. S6b**) and a slight, but non-statistically significant, increase in CD36 mRNA level (**Fig. S6c)**. FACS analysis revealed a greater than two-fold increase of CD36 cell surface expression (**Fig. 7e, f**). On the other hand, incubating EndoC-βH1 cells with recombinant PCSK9 did not affect CD36 cell surface expression (**Fig. 7g-h)**. Finally, upon PCSK9 knockdown, fatty acid uptake increased by 50% (**Fig. 7i**).

**Figure 7:**
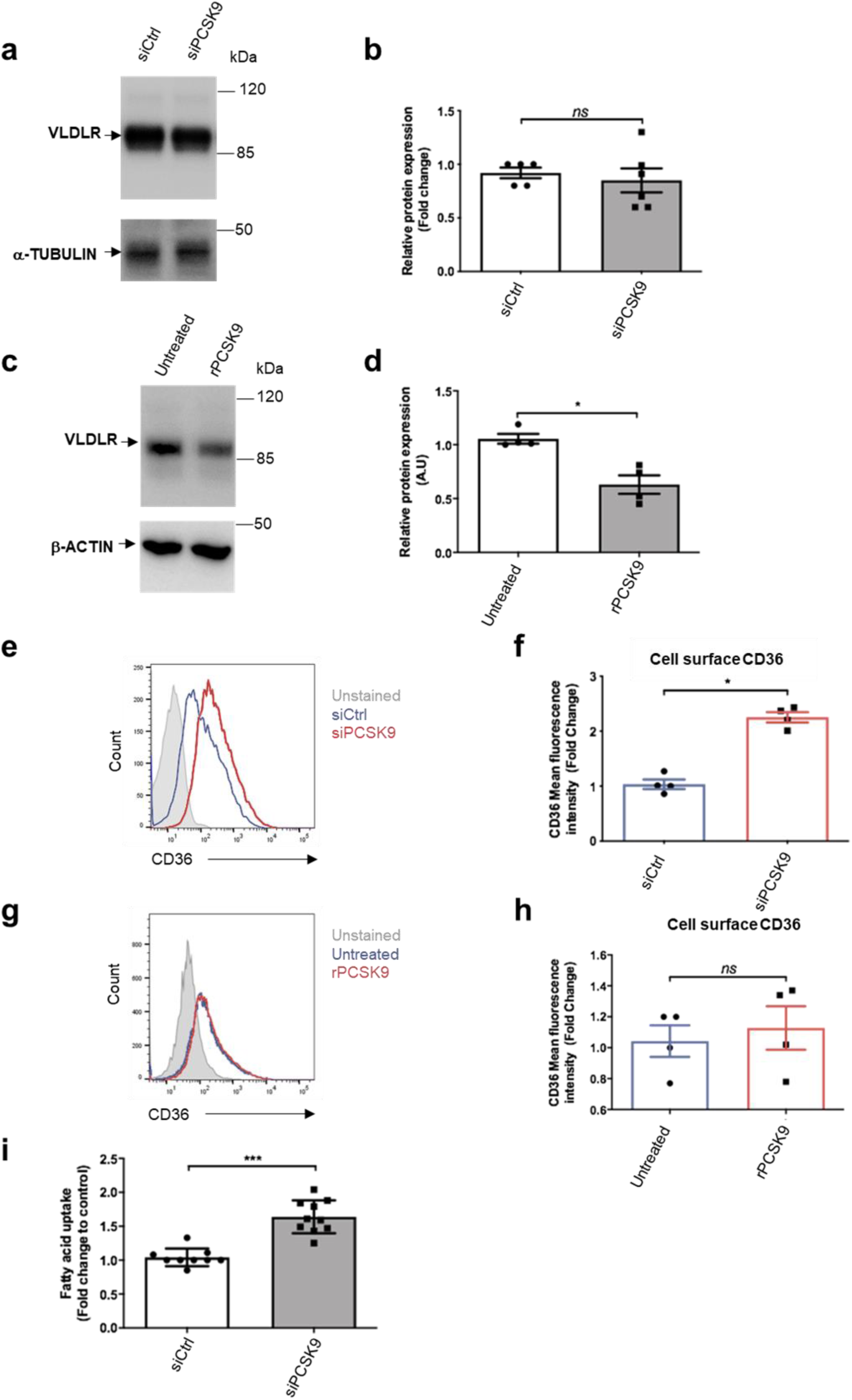
PCSK9 regulates the expression of VLDLR and CD36, 2 actors involved in fatty acid homeostasis. (**a, b**) EndoC-βH1 cells were transfected with control nontarget siRNA (siCtrl) or siRNA targeting PCSK9. Three days later, VLDLR expression was studied by Western blot and quantified *(n=5-6*). (**c, d**) EndoC-βH1 cells were exposed for 16 hours to human recombinant PCSK9 (2.5 µg/ml) and VLDLR protein expression was studied by Western blot (*n=4*). (**e, f**): EndoC-βH1 were transfected with siRNA (siCtrl) or siRNA targeting PCSK9. Six days later, cell surface CD36 was analyzed by FACS (*n=4*). (**g, h**) EndoC-βH1 cells were exposed for 16 hours to 2.5 µg/ml human recombinant PCSK9 and cell surface CD36 was analyzed by FACS (*n=4*). (**i**) Fatty acid uptake was evaluated six days following PCSK9 knockdown (*n =10*). Data represent the means ± SEM. *p < 0.05, ***p < 0.001

In conclusion, both extracellular and intracellular PCSK9 can modulate beta cell fatty acid metabolism through the regulation of VLDLR and CD36 expression respectively.

## Discussion

Here, we demonstrate that beta cells produce and secrete PCSK9 in a tightly controlled manner. We also observe that PCSK9 regulates the expression of a number of proteins, including LDLR and VLDLR through extracellular mechanisms, and CD36, HLA-ABC and PD-L1 through intracellular mechanisms.

In liver cells, PCSK9 is expressed as a pro-enzyme, processed intracellularly into its mature form and finally secreted [1]. By using EndoC-βH1 cells [33], we demonstrated here that it is also the case in human beta cells. Indeed, Western blot analyses detected both Pro-PCSK9 and mature PCSK9 in cell extracts, while only mature PCSK9 was detected in conditioned medium. We noted that the mature PCSK9/PCSK9 total ratio and secreted/intracellular PCSK9 ratio was lower in beta cells compared to liver cells. This suggest that Pro-PCSK9 processing is more efficient in liver cells compared to beta cells and that intracellular PCSK9 pool is higher in beta cells. This may explain the preferential intracellular regulation of some proteins by PCSK9. The mechanisms that regulate Pro-PCSK9 processing and secretion have only been partially characterized [4]. Our results also indicate that PCSK9 processing and secretion might be differentially regulated in liver and beta cells. Beta cells might thus be used as an additional model to study this complex processing.

In liver cells, the transcriptional regulation of PCSK9 has been studied in great detail [4]. It has been shown that SREBP2 regulates both basal and Mevastatin-induced expression of PCSK9 [18, 53, 54]. Interestingly, we demonstrate that in beta cells, physiological PCSK9 levels are regulated by SREBP1, while pharmacological regulation of PCSK9 by Mevastatin is SREBP2-dependent. This is intriguing as in the case of other SREBP1 targets such as FASN and SCD, we observe a SREBP2 mechanism that compensates for SREBP1-mediated downregulation. This compensatory mechanism is not evident for PCSK9, indicating that physiological levels of PCSK9 in beta cells are highly SREBP1-dependent.

The role of PCSK9 in pancreatic islet function is currently debated. Some studies indicate that PCSK9-deficient mice develop hyperglycemia and insulinopenia [29, 30], while others do not observe this phenotype [28, 31]. The reasons for these discrepancies might be linked to the age and/or genetic background of the mice. Here, we observed a sharp decrease in several beta cell enriched transcription factors upon PCSK9 knockdown in EndoC-βH1 cells. It is for example the case for *PDX1* and *NKX6-1* that are known to play major roles in beta cell development and function [55–57]. We also detected some perturbation in glucose-stimulated insulin secretion with a specific increase in basal insulin secretion, which was not addressed by Ramin-Mangata *et al*. [32]. Taken together, the phenotype observed upon PCSK9 knockdown (decreased expression of specific transcription factors and increased basal insulin secretion) resembles that observed in immature fetal beta cells [58, 59] and in insulin-producing beta cells differentiated *in vitro* from multipotent stem cells [60, 61]. It suggests that PCSK9 might be important to maintain a mature phenotype in pancreatic beta cells.

In mouse islets, loss of beta-cell PCSK9 results in unchanged LDLR protein levels [31]. Unexpectedly, data from a recently published study [32] indicate that PCSK9 knockdown in EndoC-βH1 cells increased LDLR cell surface expression by 20%. Our data do not support this last claim. We observed, as expected, that exogenously added PCSK9 decreased the expression of LDLR at the cell surface. On the other hand, efficient PCSK9 knockdown did not modify neither *LDLR* mRNA, nor the total amounts of LDLR protein measured either by Western blot or by global mass spectrometry, nor cell surface LDLR levels measured by FACS. Moreover, we observed similar results when looking at VLDLR, yet another PCSK9 target [9]: VLDLR protein levels decreased upon treatment with exogenous PCSK9, without any effect upon PCSK9 knockdown. Our data is consistent with observations in mouse islets [31]. The reasons for the discrepancy with Ramin-Mangata et al. [32] are difficult to discuss, as this study presents FACS data in arbitrary units without supporting FACS plots. Taken together, we propose that in pancreatic beta cells, LDLR and VLDLR levels are regulated by PCSK9 extracellularly rather than through intracellular degradation.

CD36 has been described as another PCSK9 target both in mouse liver and in adipocytes [12]. There, PCSK9 increases CD36 degradation by a mechanism involving both lysosomes and proteasomes [12]. Interestingly, in pancreatic beta cells, inhibiting CD36 has been described as a protective mechanism to prevent excessive lipid accumulation [62], while increased CD36 expression was found deleterious for proper beta cell function [63]. Here, we observe that PCSK9 knockdown in EndoC-βH1 increased cell surface CD36 expression and fatty acid uptake. Based on RNA-seq and proteomics in PCSK9 knockdown cells and the insensitivity of cell surface CD36 expression to recombinant PCSK9, we postulate that PCSK9 regulates CD36 by an intracellular mechanism. The precise mechanism of this regulation was not identified, but likely involves both proteasomal and lysosomal degradation [12]. Further experiments, in particular by using proteasome or lysosome inhibitors are needed to elucidate the nature of CD36 regulation by PCSK9 in the pancreatic beta cell.

In tumoral cells, MHC-I was recently described as a new PCSK9 target, as PCSK9 down regulates MHC-I cell surface expression by increasing its lysosomal degradation and disrupting its recycling [15]. In this model, the authors observed marginal effects of PCSK9 on the immune regulatory protein PDL-1. Interestingly, MHC-I and PDL-1 play major roles in beta cells. Their expression increases in pancreatic beta cell during inflammatory responses [49, 50, 64] and is also induced in islets from type 1 diabetic patients [50, 64]. There, it has been proposed that autoreactive CD8 T cells interact with beta cells and kill them through an MHC-I mediated activation while PD-L1 would be protective against autoimmune injuries [65]. Here, we found that PCSK9 knockdown increases both PD-L1 and HLA-ABC cell surface expression, particularly upon Interferon-*γ* stimulation. On the other hand, recombinant PCSK9 did not modulate HLA-ABC and PD-L1 levels. We thus propose that PCSK9 regulates the expression of PD-L1 and HLA-ABC in human beta cells by an intracellular mechanism.

In conclusion, we find that PCSK9 is expressed and secreted by human pancreatic beta cells in a tightly regulated manner, which is altered in the presence of statins. In turn, we find that PCSK9 regulates cell surface expression of CD36, PD-L1 and HLA-ABC through intracellular mechanisms, but regulates LDLR and VLDLR expression through an extracellular mechanism in pancreatic beta cells. It will be interesting to determine if PCSK9 monoclonal antibodies or siRNA-based therapies against PCSK9 affect expression levels of LDLR and VLDLR, or CD36, PD-L1, and HLA-ABC respectively in pancreatic beta cells, and whether co-treatment with statins interferes with the effect of PCSK9 siRNA in the pancreas.

## Data availability

The datasets generated during and/or analyzed during the current study are available from the corresponding authors.

Data sets accession numbers: RNA-seq GSE182016, cell lysate TMT proteomics experiment PXD027921, cell lysate DIA experiment PXD027911, secretome DIA proteomics PXD027913.

## Author contribution statements

K.S., G.H., I.G. and C.B. performed experiments. M.A. and F.C. provided human islets. K.S., R.S., M.R. and A.J. analyzed data and drafted the manuscript with contributions from S.A., W.L.G., K.SG, C.R.U.

All authors approved the final draft.

## Acknowledgments

The authors thank Natascha de Graaf (LUMC) for technical support.

The RS laboratory is supported by grants from the Fondation pour la Recherche Médicale (EQU201903007793), the Dutch Diabetes Research Foundation, the DON Foundation, the Fondation Francophone pour la Recherche sur le Diabetes (FFRD), the Agence Nationale de la Recherche (ANR-19-CE15-0014-01), Laboratoire d’Excellence consortium Revive, the Innovative Medicines Initiative 2 Joint Undertaking under grant agreement no. 115797 (INNODIA) and no. 945268 (INNODIA HARVEST). This joint undertaking receives support from the Union’s Horizon 2020 research and innovation program EFPIA, JDRF, and the Leona M. and Harry B. Helmsley Charitable Trust.

