## Supplemental data for "PCSK9 affects expression of key surface proteins in human pancreatic beta cells through intra- and extracellular regulatory circuits"

### Supplementary Table 1: Human islets donors characteristics

#### Checklist for reporting human islet preparations used in research

Adapted from Hart NJ, Powers AC (2018) Progress, challenges, and suggestions for using human islets to understand islet biology and human diabetes. Diabetologia https://doi.org/10.1007/s00125-018-4772-2

| Islet preparation | 1 | 2 | 3 | 4 | 5 | 6 | 7 | 8 | 9 | 10 | 11 |
|---|---|---|---|---|---|---|---|---|---|---|---|
| | | | | | | | | MANDATORY INFORMATION | | | |
| Unique identifier | HSL P121 | HSL P135 | HSL P137 | HSL P138 | R174 | R175 | R177 | P164 | P165 | P146 | P126 |
| Donor age (years) | 57 | 28 | 30 | 27 | 47 | 42 | 47 | 58 | 36 | 53 | 61 |
| Donor sex (M/F) | F | M | M | F | F | M | F | F | M | F | M |
| Donor BMI (kg/m$^2$) | 23.4 | 28.4 | 30.5 | 27.7 | 28 | 28 | 26 | 29 | 25.5 | 29.9 | 23.8 |
| Donor $HbA_{1c}$ or other measure of blood glucose control | NT | NT | NT | NT | 7,2mM (last glucose measurement at ICU) | - | 6,0 mM (last glucose measurement at ICU) | | | | |
| Origin/source of islets[b] | APHP/hospital St Louis, Paris | APHP/hospital St Louis, Paris | APHP/hospital St Louis, Paris | APHP/hospital St Louis, Paris | Leiden University Medical Center | Leiden University Medical Center | Leiden University Medical Center | APHP/hospital St Louis, Paris | APHP/hospital St Louis, Paris | APHP/hospital St Louis, Paris | APHP/hospital St Louis, Paris |
| Islet isolation centre | Saint-Louis hospital | Saint-Louis hospital | Saint-Louis hospital | Saint-Louis hospital | Leiden University Medical Center | Leiden University Medical Center | Leiden University Medical Center | Saint-Louis hospital | Saint-Louis hospital | Saint-Louis hospital | Saint-Louis hospital |
| Donor history of diabetes? Please select yes/no from drop down list | No | No | No | No | No | No | No | No | No | No | No |
| | | | | | | | If Yes, complete the next two lines if this information is available | | | | |
| Diabetes duration (years) | | | | | | | | | | | |
| Glucose-lowering therapy at time of death[c] | | | | | | | | | | | |
| | | | | | | | | RECOMMENDED INFORMATION | | | |
| Donor cause of death | Vascular | Trauma | Vascular | Vascular | CVA | CVA | Respiratory | Vascular | Vascular | Vascular | Vascular |
| Warm ischaemia time (h) | | | | | 17 min | 19 min | 22 min | | | | |
| Cold ischaemia time (h) | 8:09 | 9:00 | 07:45 | 09:07 | 21:40 | 16:43 | 18:02 | | | | |
| Estimated purity (%) | 70 | 70 | 80 | 80 | 99 | 99 | 95 | 80-100 | 80-100 | 57 | 80 |
| Estimated viability (%) | NT | NT | NT | NT | NT | NT | NT | NT | NT | NT | NT |
| Total culture time (h)[d] | 168 | 168 | 168 | 132 | | | | | | | |
| Glucose-stimulated insulin secretion or other functional measurement[e] | No | Yes | Yes | Yes | | | | | | | |
| Handpicked to purity? Please select yes/no from drop down list | Yes | Yes | Yes | Yes | | | | | | | |
| Additional notes | Fig. 1b | Fig. 1b | Fig. 1b | Fig. 1b | Fig. 1b, Fig. 1h | Fig. 1b, Fig. 1h | Fig. 1b, Fig. 1h | Fig. 1h | Fig. 1h | Fig. 1h | Fig. 1b, Fig. 1h |

[a]If you have used more than eight islet preparations, please complete additional forms as necessary
[b]For example, IIDP, ECIT, Alberta IsletCore
[c]Please specify the therapy/therapies
[d]Time of islet culture at the isolation centre, during shipment and at the receiving laboratory
[e]Please specify the test and the results

### Supplementary Table 2: List of primer used for RT-qPCR

#### Human primers

| Gene | Forward primer | Reverse primer |
|---|---|---|
| *CYCLOPHILIN* | ATGGCAAATGCTGGACCCAACA | ACATGCTTGCCATCCAACCACT |
| *PCSK9* | TCCACGCTTCCTGCTGCCAT | CAGGCAGTCAGGGTCCAGCC |
| *LDLR* | GAGGTCCACATTTGCGACAAC | GTCATCCTCCAGACTGACCATC |
| *HMGCR* | ATTTTGGGTATTGCTGGCCTT | ATGTGCTTGCTCTGGAAAGGT |
| *FASN* | TACGACTACGGCCCTCATTT | CCATGAAGCTCACCCAGTTATC |
| *SCD* | ACAACTACCACCACTCCTTTC | GGAGACTTTCTTCCGGTCATAG |
| *SREBF-1* | GCACCCACTCCATTGAAGAT | GGCACTGACTCTTCCTTGATAC |
| *SREBF-2* | ATGGGCAGCAGAGTT | CGACAGTAGCAGGTC |
| *CASR* | GCTCTTCACCAATGGCTCCTGT | CCACACTCATCAAAGGTCACCTG |
| *APP* | CCTTCTCGTTCCTGACAAGTGC | GGCAGCAACATGCCGTAGTCAT |
| *APLP-2* | GATCGGTGCCGAAGAGAAAG | CCACGGATTCCCGCTCTT |
| *MDK* | GCTACAATGCTCAGTGCCAGGA | CTTGGCGTCTAGTCCTTTCCCT |
| *SCG2* | GAGAAGCCGAATGGATCAGTGG | TCTGGATGGTCTAAGTCAGCCTC |
| *FADS1* | TCATCAGCCACTACGCCG | GCTCTTTATTCTTGGTGGGCTC |
| *SCD5* | GGTGGCTGTTTGTTCGCAAG | ACCACAAAGCACATGAGCAC |
| *VLDLR* | CAAGGATGGCAGTGATGAGGTC | CTCGGATACCATTACACTGCCTG |
| *CD36* | CAGGTCAACCTATTGGTCAAGCC | GCCTTCTCATCACCAATGGTCC |
| *PD-L1* | CAGCCGCGCTTCTGT | ACATATAGGTCCTTG |
| *CMH-I* | GAGAACGGGAAGGAG | CATCTCAGGGTGAGG |
| *PDX-1* | TACTGGATTGGCGTTGTTTGTGGC | AGGGAGCCTTCCAATGTGTATGGT |
| *NKX6-1* | GCCCTGGGAGAAGACTTTCGAACAA | TTCTGGAACAGACCTTGACCTGA |

#### Mouse primers

| Gene | Forward primer | Reverse primer |
|---|---|---|
| Cyclophilin | CAGGTCCTGGCATCT | TGCTTGCTGGTCTTG |
| Ins-1 | CAGAGACCATCAGCAAGCAG | GGGACCACAAAGATGCTGTT |
| Gcg | TGAAGACAAACGCCACTCAC | TGACGTTTGGCAATGTTGTT |
| Sst | TCCGTCAGTTTCTGCAGAAGTCTC | GTACTTGGCCAGTTCCTGTTTCCC |
| Pcsk9 | ACGCCTGCCTCTACTCCCCA | GTAACTGGCTGGTCCTGGGC |

Figure S1

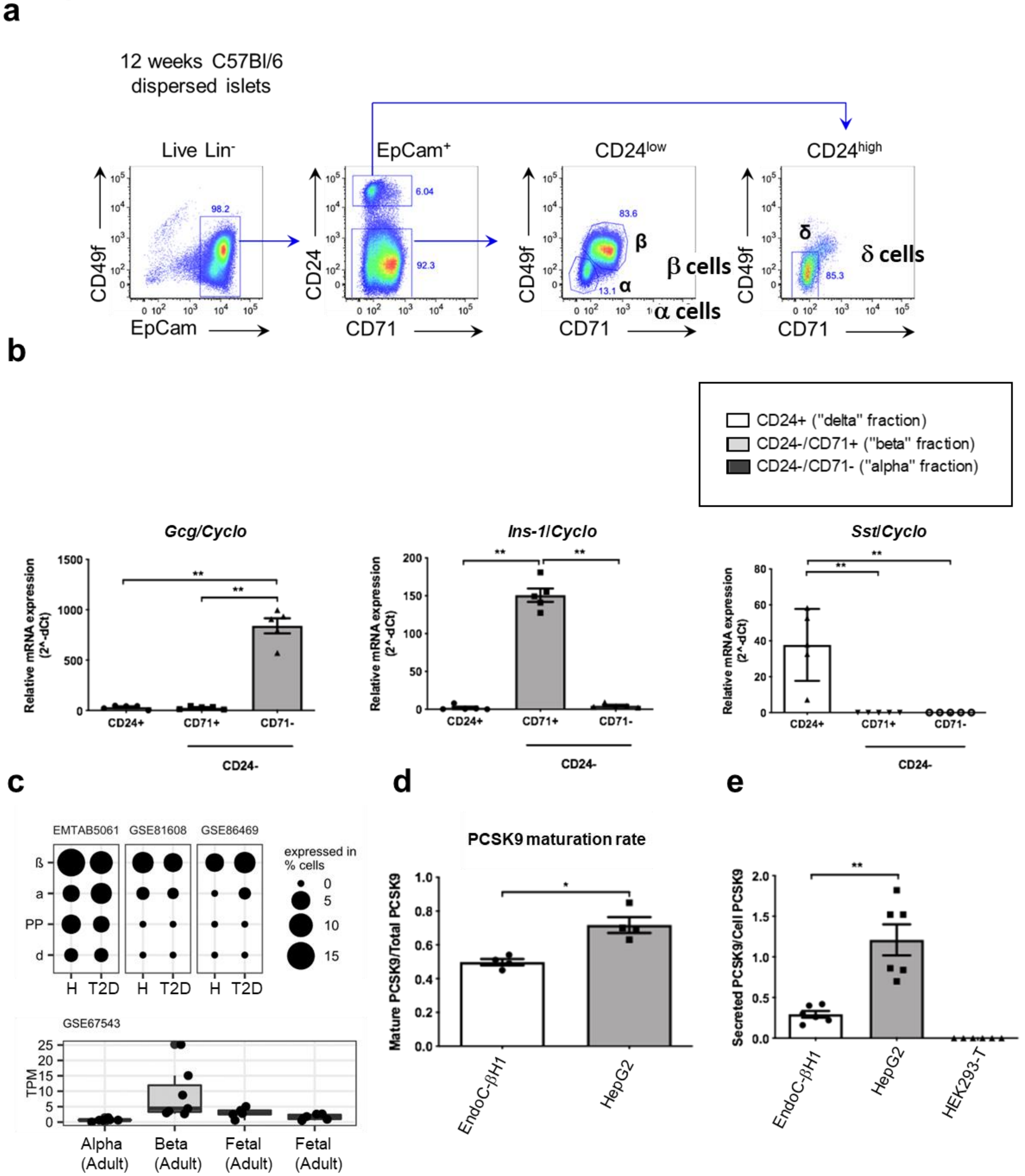


(**a**) Representative flow cytometry plots depicting the FACS-based strategy adopted to purify alpha ($CD71^-$), beta ($CD71^+$) and delta ($CD24^+$) cells enriched populations. (**b**) RT-qPCR analysis of glucagon (*Gcg*), insulin-1 (*Ins-1*) and somatostatin (*Sst*) in alpha, beta and delta cell enriched populations (*n=5*). (**c**) PCSK9 transcript levels measured by RNA-seq from healthy (H), type 2 diabetes (T2D) human pancreatic islets (EMTAB5061, GSE81608, GSE86469), and in alpha and beta cell populations sorted by FACS from adults and fetal human islets (GSE67543). (**d**) PCSK9 maturation rate measured as the mature/total intracellular PCSK9 ratio in EndoC-βH1 and HepG2 cells (*n=4*). (**e**) PCSK9 in conditioned media from EndoC-βH1 and HepG2 cells, normalized to PCSK9 cell content (*n=6*). Data represent the means ± SEM. *$p < 0.05$, **$p < 0.01$

### Figure S2

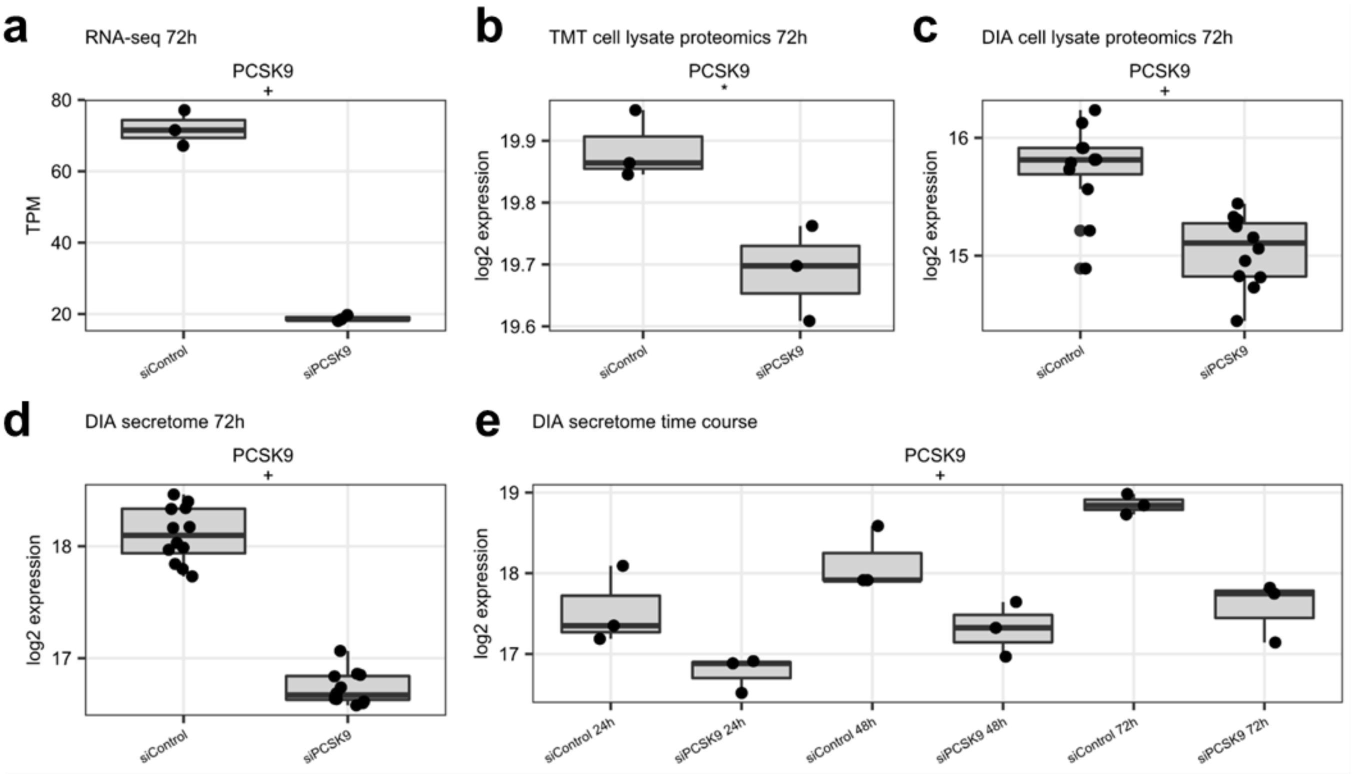


Boxplots indicating PCSK9 levels in the omics experiments. Data represent the median expression and interquartile ranges. + FDR < 0.05 (genome/proteome-wide significant), and * a raw p < 0.05.

### Figure S3

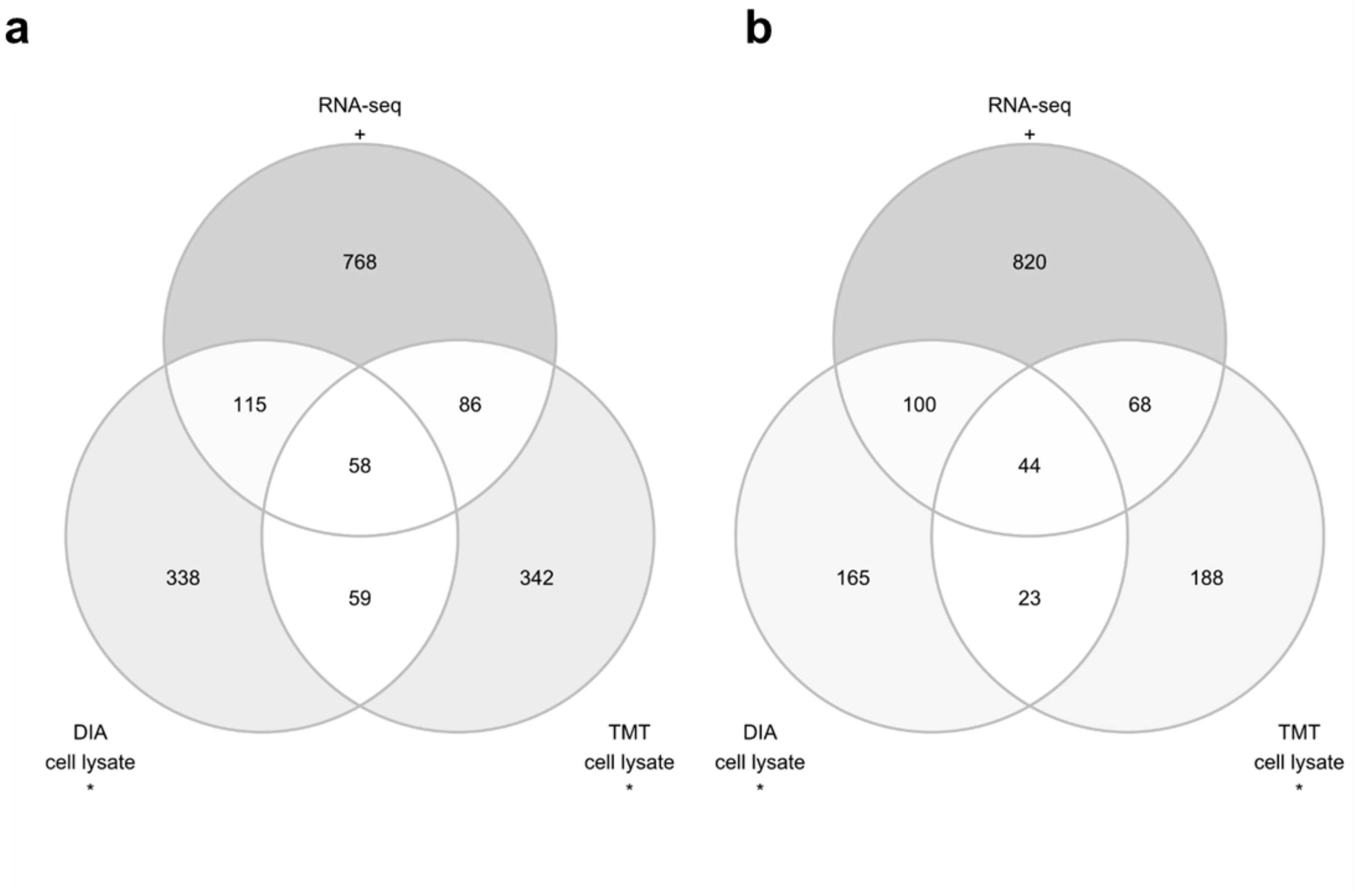


Venn diagrams illustrating agreement between the RNA-seq and proteomics experiments. (**a**) Genes and proteins up-regulated in siPCSK9 vs siControl EndoC-βH1 cells. (**b**) Genes and proteins down-regulated in siPCSK9 vs siControl EndoC-βH1 cells. Different stringency cut-off is applied for RNA-seq vs proteomics data in this figure. Genes with FDR < 0.05 are counted, while proteins are counted if they reach at least nominal significance (raw p < 0.05).

### Figure S4

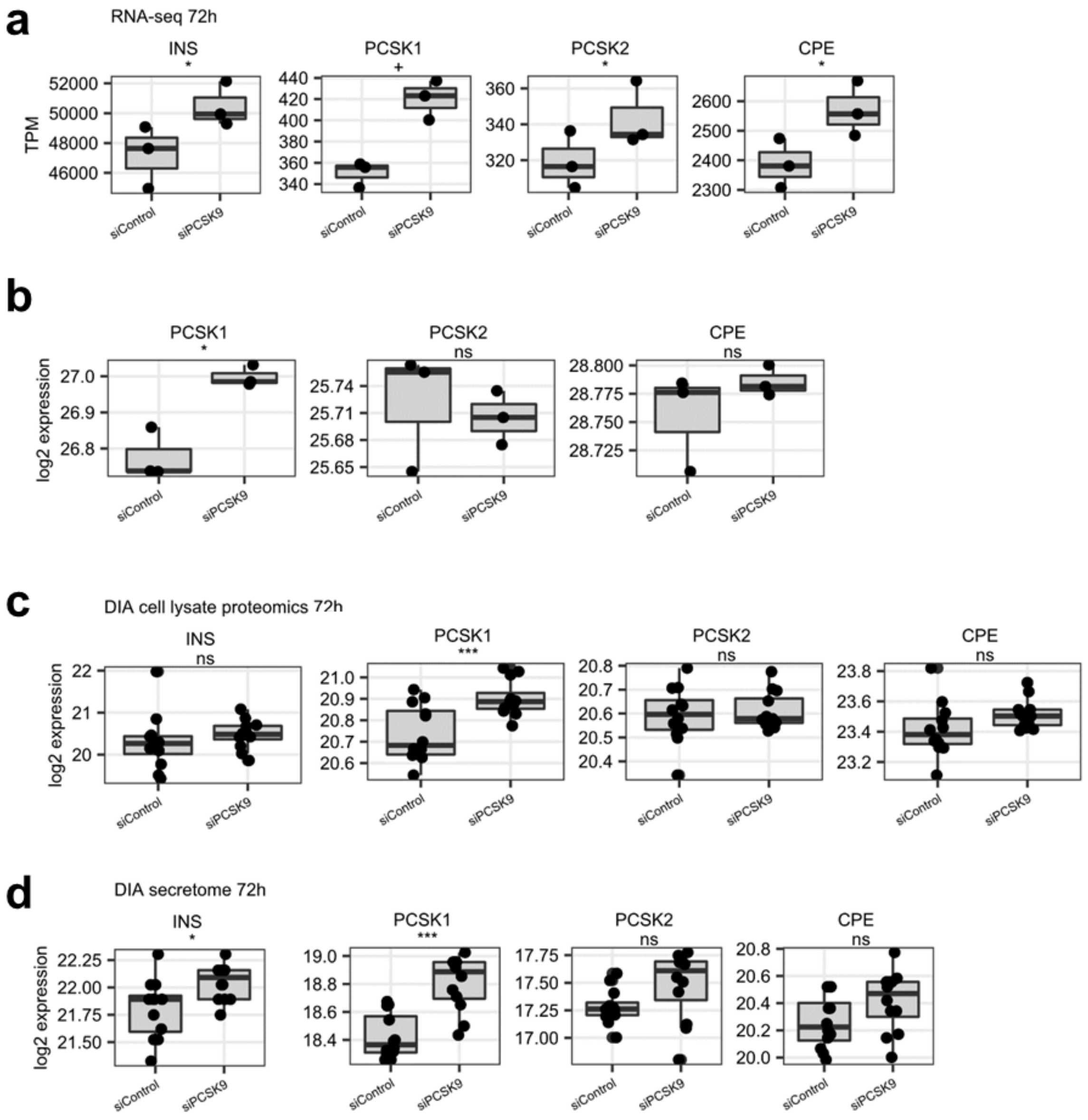


Boxplots indicating insulin (INS) and proteins involved in pro-insulin processing in (**a)** RNA-seq, (**b**) TMT cell lysate proteomic, (**c**) DIA cell proteomic and (**d**) DIA secretome experiments. Data represent the median expression and the interquartile ranges. + indicate an FDR < 0.05 (genome/proteome-wide significant), and “*”, p < 0.05, “**”,p < 0.01 and “***, p< 0.001.

Figure S5

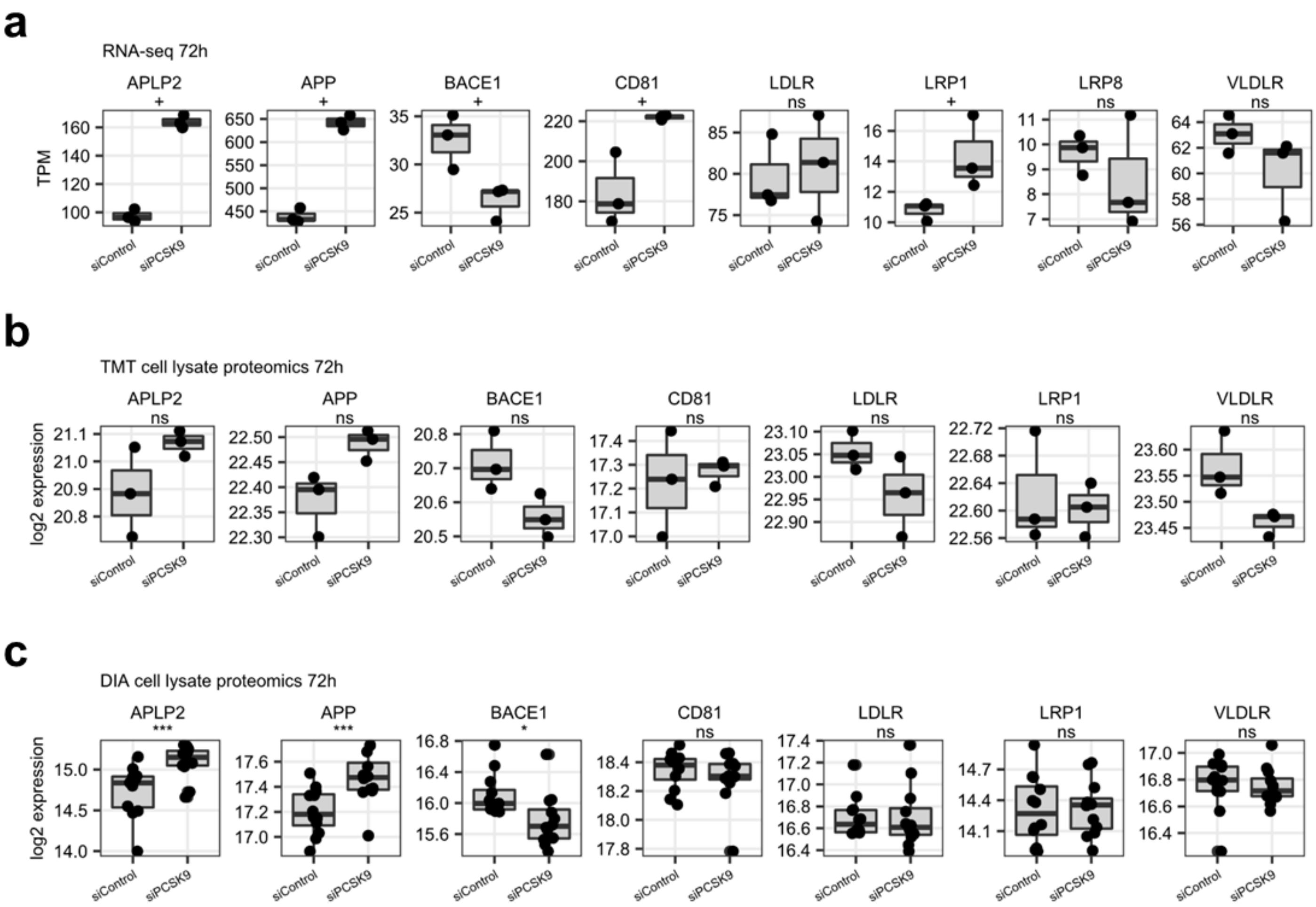


Boxplots indicating the expression of known PCSK9 targets in (**a**) RNA-seq, (**b**) TMT cell lysate proteomic, (**c**) DIA cell proteomic. Data represent the median expression and the interquartile ranges. + indicate an FDR < 0.05 (genome/proteome-wide significant), and “*”, p < 0.05, “**”,p < 0.01 and “***, p< 0.001.

### Figure S6

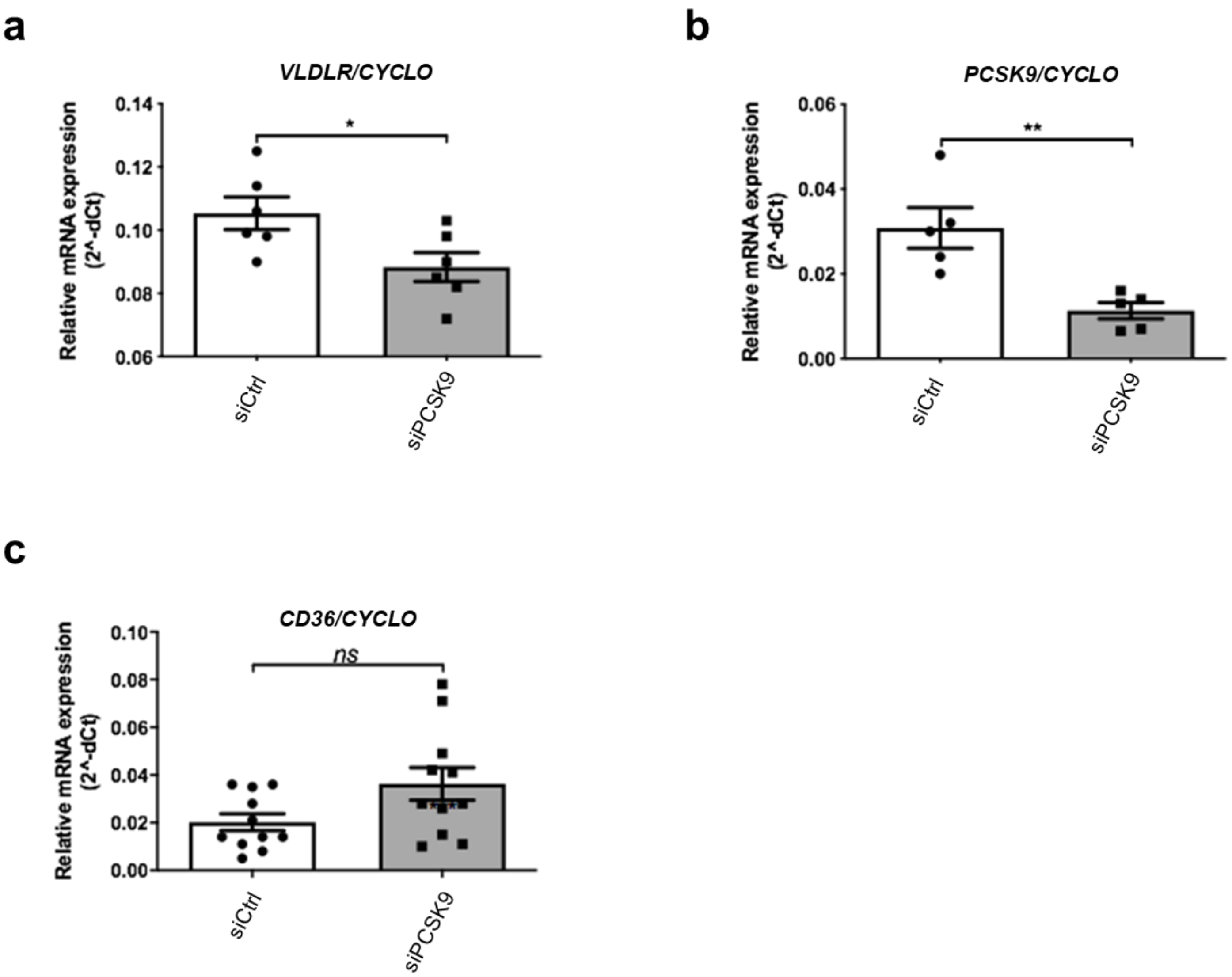


EndoC-βH1 cells were transfected with control nontarget siRNA (siCtrl) or siRNA targeting PCSK9. Three days later, VLDLR expression was studied by RT-qPCR *(n=6)* (**a**). Six days later, PCSK9 (*n=5*) (**b**) and CD36 (*n=11*) (**c**) expressions were analyzed by RT-qPCR.
